## Supplemental Figures for "Multiple time-scales of decision making in the hippocampus and prefrontal cortex"

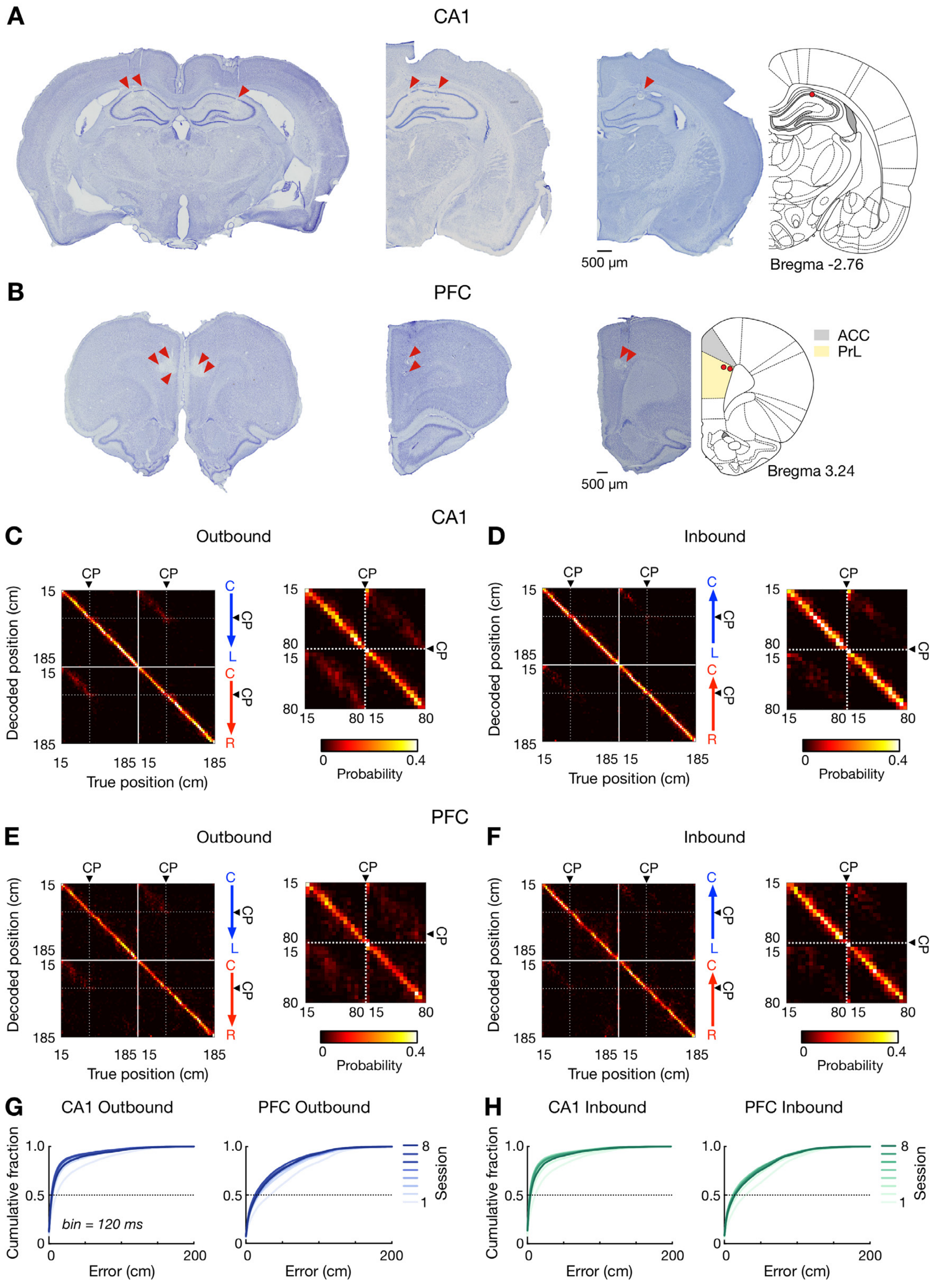

**Figure S1. Recording locations and behavioral-sequence representations of locations over learning**

**(A** and **B)** Representative Nissl-stained coronal sections showing tetrode positions in **(A)** CA1 and **(B)** PFC (primarily in PreLimbic, PrL, cortex). Lesion locations at the end of a tetrode track are indicated by red arrowheads. *Left*: sections from one animal with 64 tetrodes targeting the bilateral CA1 of dorsal hippocampus and PFC. *Middle* and *Right*: sections from animals with 32 tetrodes implanted over the right hemisphere. The brain sections on the right are mapped onto a stereotaxic atlas (Paxinos and Watson, 2004) (the distance from Bregma in mm denoted).

**(C-F)** Confusion matrices depicting actual (*x*-axis) and decoded (*y*-axis) positions in **(C** and **D)** CA1 and **(E** and **F)** PFC. For each plot pair, decoding along entire trajectories is shown on the left, and the decoding within the center stem is enlarged on the right.

**(G** and **H)** Cumulative position decoding errors (bin = 120 ms) across all sessions for **(G)** outbound and **(H)** inbound. Each line represents a single session (color coded). Locations within 15 cm of the reward well were excluded to prevent contamination from SWR activity. Overall decoding errors: CA1 outbound, 6.00 ± 0.08 cm; CA1 inbound, 6.00 ± 0.07 cm; PFC outbound, 18.00 ± 0.10 cm; PFC inbound, 16.00 ± 0.09 cm (median ± SEM).

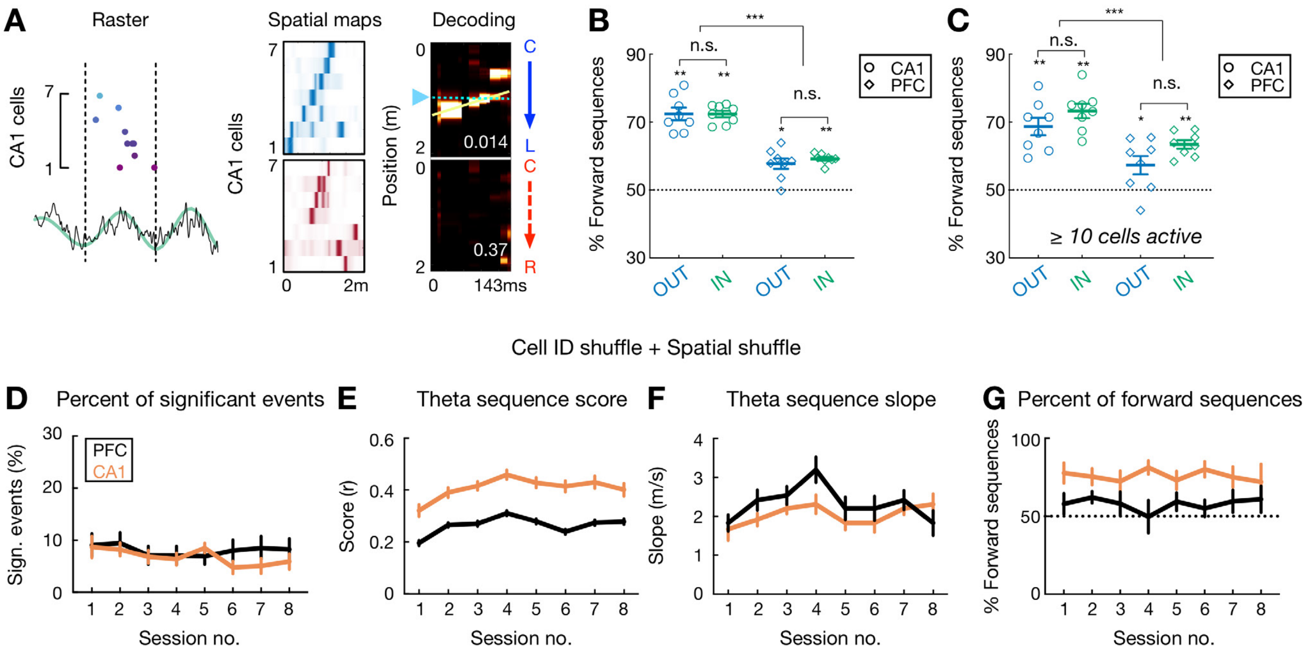


**Figure S2. Detection of theta sequences in CA1 and PFC is statistically robust**

**(A)** An example of reverse theta sequences in CA1. Data are presented as in **Figure 2A**.

**(B)** Percent of forward theta sequences during outbound (OUT; blue) and inbound (IN; green) running. Consistent with previous reports in CA1 (Drieu et al., 2018; Gupta et al., 2012; Wikenheiser and Redish, 2015; Zheng et al., 2016), theta sequences in CA1 and PFC are overall biased toward a forward order (**$p$ = 0.0078, *$p$ = 0.0156, signed-rank tests compared to 50%), and this bias is stronger in CA1 than PFC (***$p$ = 0.0002, Friedman test with Dunn's *post hoc*). Each symbol represents a session, summing over all 9 animals. Error bars: mean ± SEM.

**(C)** Percent of forward theta sequences using a more stringent criterion of at least 10 cells active per theta cycle. Data are presented as in **(B)**. Note that similar proportions of forward versus reverse sequences were detected independent of the cell thresholds (compared with **B**).

**(D-G)** Significant theta sequences are detected in both CA1 (orange lines) and PFC (black lines) by applying additional criteria for significance in additional to the spatial shuffle (a shuffle *post* the Bayesian decoding; see **METHODS**). With an additional cell-identity shuffle (a shuffle *prior to* the Bayesian decoding), a significant theta sequence would additionally belong to the top 95$^{th}$ percentile of the shuffled distribution of goodness-of-fits ($R_{max}$), and exceed the 97.5$^{th}$ percentile or be below the 2.5$^{th}$ percentile (for reverse sequences) of the shuffled distribution of weighted correlations. **(D)** Approximately 50% of all theta sequences in both CA1 and PFC met these additional stringent criteria, and the proportions are similar in CA1 and PFC ($p$ = 0.11, session-by-session rank-sum paired tests) regardless of the shuffling methods (compared with **Figure 2G**). **(E)** Theta sequence score, **(F)** Theta sequences slope, and **(G)** Percent of forward theta sequences with the additional criteria for significance. Data are presented as in **Figures 2G-2J**.

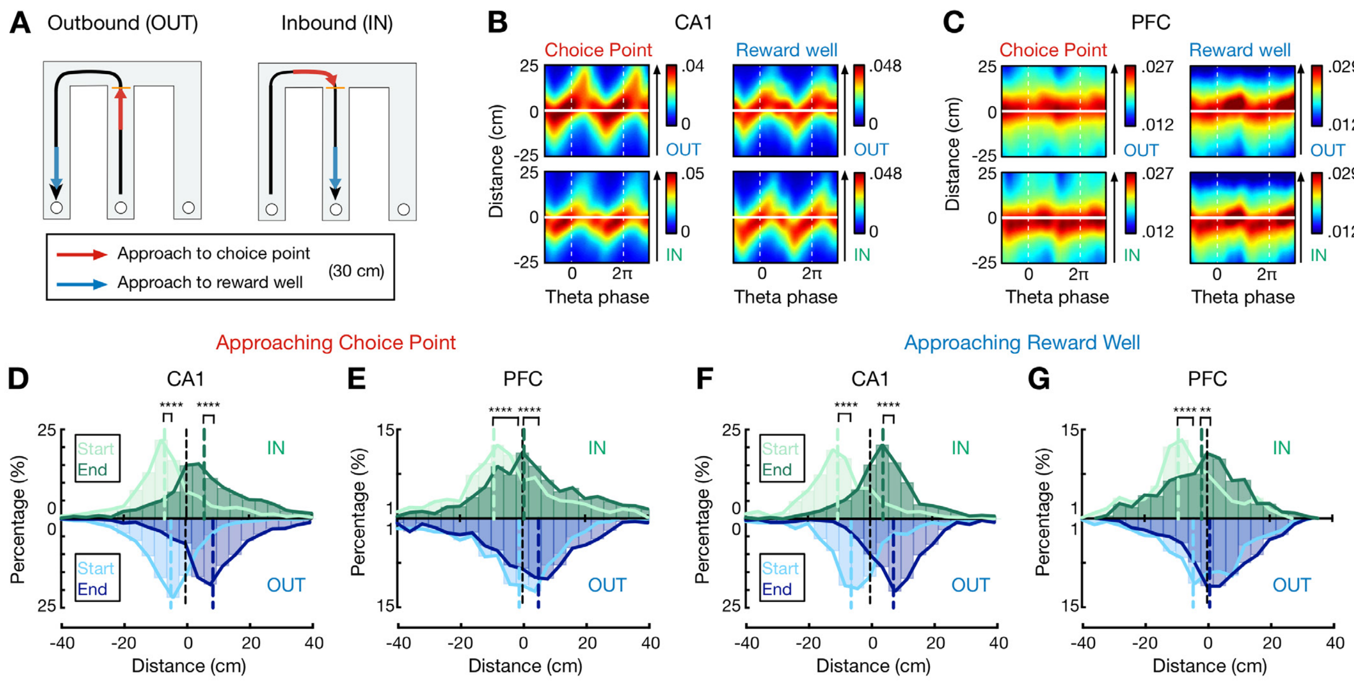


**Figure S3. Look-ahead of theta sequences was similar during approach to choice point and approach to reward well**

**(A)** Schematics showing trajectory segments used for approach to choice point (red) and approach to reward well (blue) during outbound (*left*) and inbound (*right*) passes. The trajectory segment used for approach to choice point was defined as the segment 30 cm before reaching the choice point, and the trajectory segment used for approach to reward well was defined as the one 30 cm before reaching the reward-well location (i.e., 45-15 cm relative to the reward well, as the locations within 15 cm of the reward well were excluded to prevent contamination from SWR activity).

**(B** and **C)** Averaged Bayesian reconstruction of all forward candidate theta sequences in **(B)** CA1 and **(C)** PFC for approach to choice point (*left*) versus reward well (*right*). Data are presented as in **Figures 3A** and **3B**.

**(D** and **E)** Distributions for start and end of reconstructed trajectories of all significant forward theta sequences in **(D)** CA1 and **(E)** PFC during approach to choice point. Data are presented as in **Figures 3C** and **3D**. ****$p$ < 1e-4, Kolmogorov-Smirnov test.

**(F** and **G)** Distributions for start and end of reconstructed trajectories of all significant forward theta sequences in **(F)** CA1 and **(G)** PFC during approach to reward well. Data are presented as in **Figures 3C**

and **3D**. ****$p$ < 1e-4, **$p$ = 0.0044, Kolmogorov-Smirnov test. Note that similar look-ahead properties of theta sequences were observed for approach to choice point versus reward well.

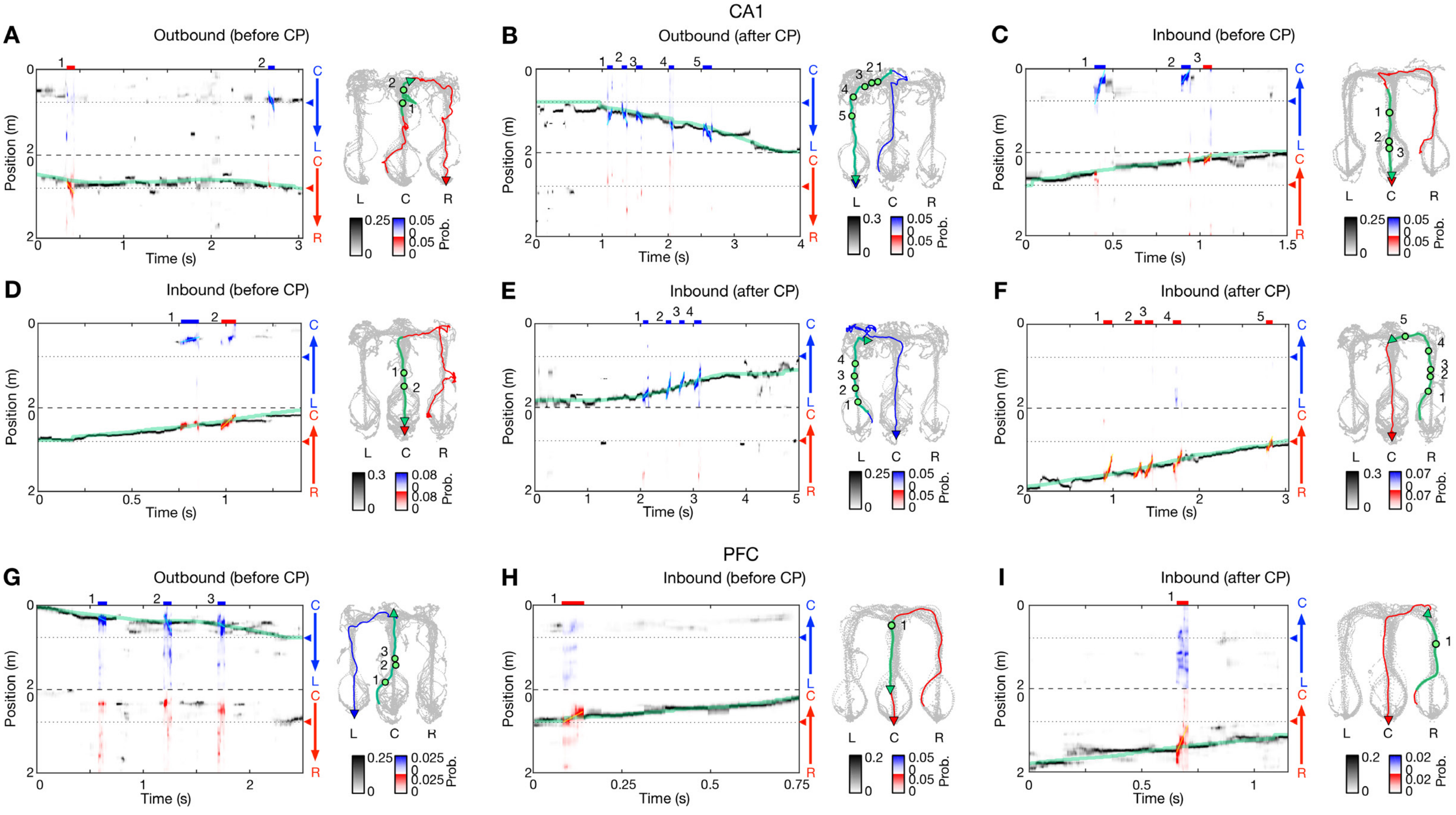


**Figure S4. Additional examples of theta-sequence representations of behavioral choices**

**(A-F)** Six additional decoding examples in CA1. **(A and B)** for outbound navigation. **(C-F)** for inbound navigation.

**(G-I)** Three additional decoding examples in PFC. **(G)** for outbound navigation. **(H and I)** for inbound navigation.

Data are presented as in **Figures 6C-6F**.

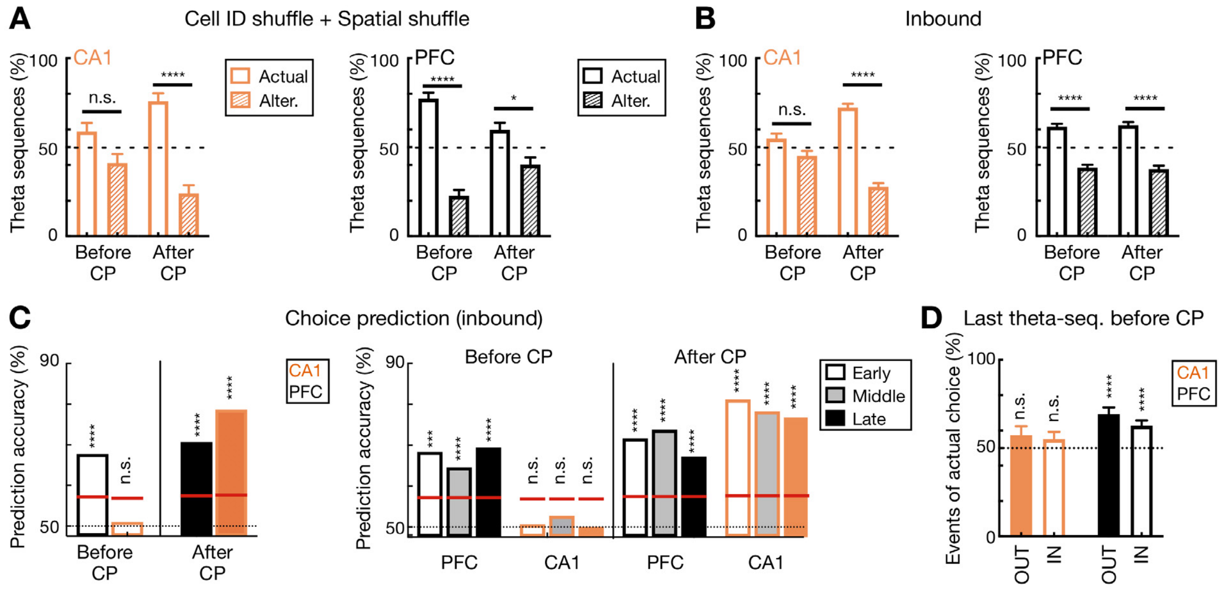


**Figure S5. Theta-sequence coding for behavioral choices during inbound navigation and additional controls**

**(A)** Percent of theta sequences representing current actual or alternative choice before (left two bars) and after CP (right two bars) during outbound navigation with the two shuffling procedures shown in **Figure S2** (****$p$ < 1e-4, *$p$ < 0.05, n.s. $p$ > 0.05, session-by-session rank-sum paired tests).

**(B)** Percent of theta sequences representing current actual or alternative choice before and after CP during inbound navigation (****$p$ < 1e-4, n.s. $p$ > 0.05, session-by-session rank-sum paired tests). Similar to that during outbound navigation (**Figure 6G**), during inbound navigation, CA1 theta sequences encoded two possible past choices on the center stem (i.e., before CP; $p$ = 0.0934, rank-sum test compared to 50%), whereas PFC theta sequences preferentially encoded the actual choice ($p$ < 1e-4).

**(C)** Trial-by-trial theta-sequence prediction of behavioral choice during inbound navigation. *Left*: During inbound navigation on the center stem, the content of CA1 theta sequences did not reliably predict animal's current choice (n.s., $p$ = 0.473, trial-label permutation tests), but the content of PFC theta sequences predicted choice well above chance (****$p$ < 0.0001). Data are presented as in **Figure 6H**. *Right*: Theta-sequence prediction of behavioral choice persists over sessions during inbound navigation (****$p$ < 0.0001, ***$p$ < 0.001, n.s, $p$ > 0.05, trial-label permutation tests). Data are presented as in **Figure 6I**.

**(D)** Choice representations by the last theta sequence before CP during outbound (OUT) and inbound (IN) navigation. The last sequence before CP of each trial was taken from the last event before exiting center stem (defined by CP) during outbound running (**Figure 1A**, *right*) and the first event after entering center stem during inbound running (**Figure 1A**, *left*). The proportions of the last sequences representing actual choices for each session are shown as mean and SEM. Solid bars are for outbound (OUT), and hollow bars are for inbound (IN). Note that the content of the last theta sequence in CA1 represented animal's current and alternative choice equivalently (orange bars; $p$ = 0.18 and 0.13 for outbound and inbound, respectively), whereas the last theta sequence in PFC preferentially represented the actual choice (black bars; $p$*'s* < 1e-4 for outbound and inbound, signed-rank test compared to 50%).

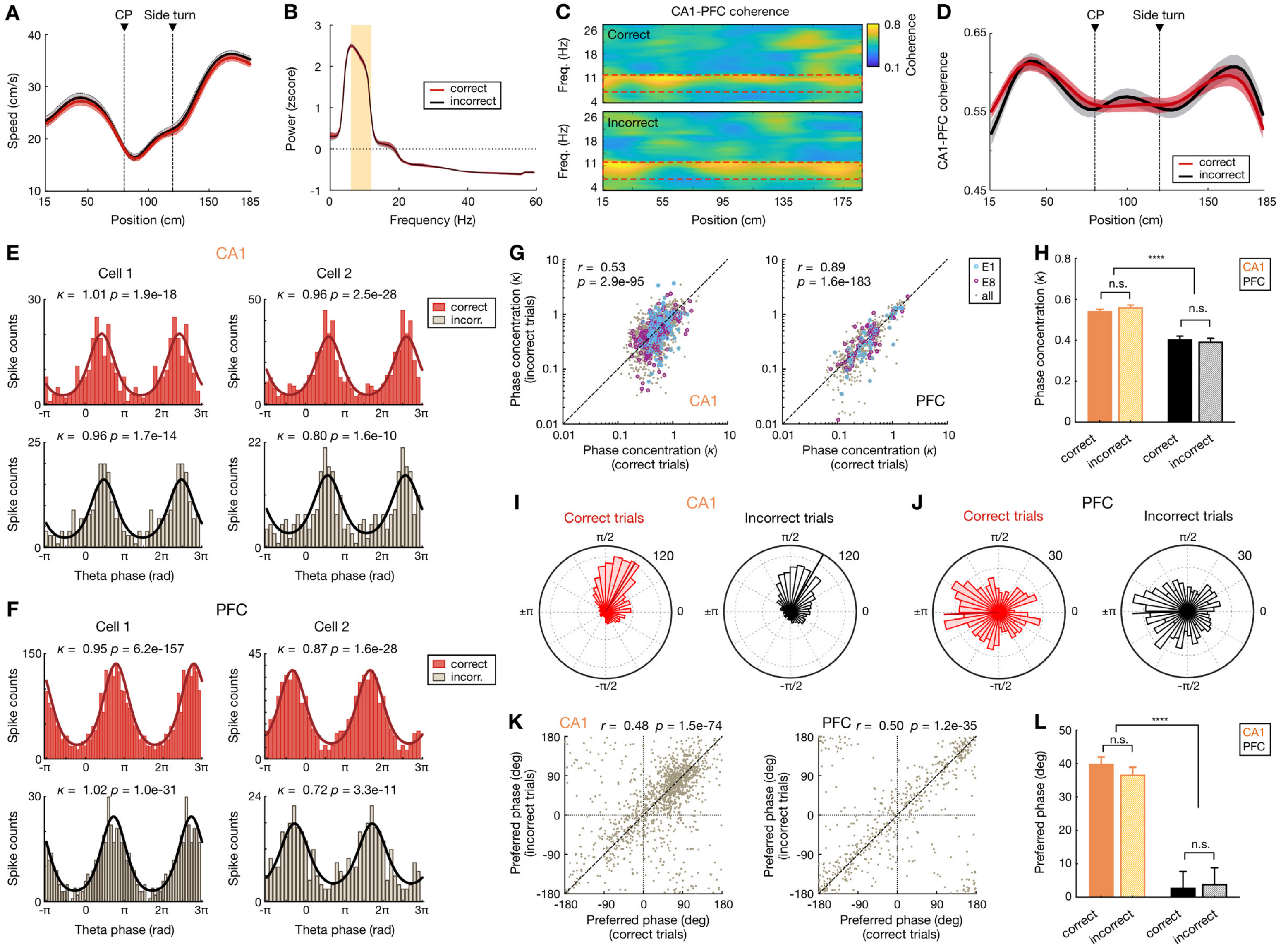


**Figure S6. Speed, theta power, coherence, and phase-locking during correct versus incorrect outbound trials**

**(A)** Animal's running speed during correct (red) versus incorrect (black) trials. Running speed was not significantly different across locations during correct versus incorrect trials (all *p's* > 0.05, rank-sum tests). Data are presented as mean and SEM.

**(B)** Power spectra during correct (red) versus incorrect (black) trials. Orange shading highlights the theta frequency range (6-12 Hz). Theta power was not significantly different during correct versus incorrect trials ($p$ = 0.79, rank-sum tests). Data are presented as mean and SD (small error bars may not be discernable in some cases).

**(C)** Averaged CA1-PFC coherograms during correct (*top*) and incorrect (*bottom*) trials of an example animal in a given session. Red boxes highlight the theta frequency range.

**(D)** CA1-PFC theta coherence was not significant different across locations during correct versus incorrect trials (all *p's* > 0.05, rank-sum tests). Data are presented as mean and SEM.

**(E** and **F)** Examples of phase locking to hippocampal theta oscillations for **(E)** two CA1 and **(F)** two PFC cells during correct (*top*) and incorrect (*bottom*) outbound trials. For each cell, spikes during all correct and incorrect trials of an example session are shown. Lines on the histograms are derived from von Mises fits (Jadhav et al., 2016; Siapas et al., 2005). Phase concentration parameter ($\kappa$) and *p*-value from Rayleigh tests are denoted. Note that while there were often fewer incorrect trials than correct trials within a given session (yielding lower spike counts for the incorrect condition), the phase-locking properties of a given cell are similar during correct versus incorrect trials.

**(G)** Phase concentration ($\kappa$) of all significantly phase-locked cells in CA1 (*left*) and PFC (*right*) during correct versus incorrect trials. Each dot represents a single cell in a given session. Blue and purple circles highlight phase concentration during the first (E1) and last (E8) sessions, respectively. Note that phase concentration is highly correlated during correct versus incorrect trials (*r*- and *p*-values denoted on the upper left corner of each plot, Pearson correlation).

**(H)** While CA1 population exhibited more concentrated phase distributions than PFC cells (*****p* < 1e-4, Kruskal-Wallis tests with Dunn's *post hoc*), consistent with previous reports (Jadhav et al., 2016; Siapas et al., 2005), phase concentration ($\kappa$) in each region is similar during correct versus incorrect trials (n.s., *p* > 0.99, Kruskal-Wallis tests with Dunn's *post hoc*).

**(I** and **J)** Polar plots of the distributions of preferred phases of all significantly phase-locked cells in **(I)** CA1 and **(J)** PFC during correct versus incorrect trials. The preferred phase of each cell was calculated from von Mises fits. Numbers on top right indicate radius of the plot (i.e., number of neurons). Note that CA1 cells preferred the falling phase of theta, while PFC cells were most often phase locked to the trough of theta, consistent with previous studies (Jadhav et al., 2016; Jones and Wilson, 2005b; Siapas et al., 2005).

**(K)** Preferred phase during correct versus incorrect trials in CA1 (*left*) and PFC (*right*). Each dot represents a single cell in a given session. Preferred phase is highly correlated during correct versus incorrect trials (*r*- and *p*-values denoted on the top of each plot, Pearson correlation).

**(L)** Distributions of preferred phase during correct versus incorrect trials (****$p$ < 1e-4, n.s., $p$ > 0.99, Kruskal-Wallis tests with Dunn's *post hoc*). Error bars: SEMs.

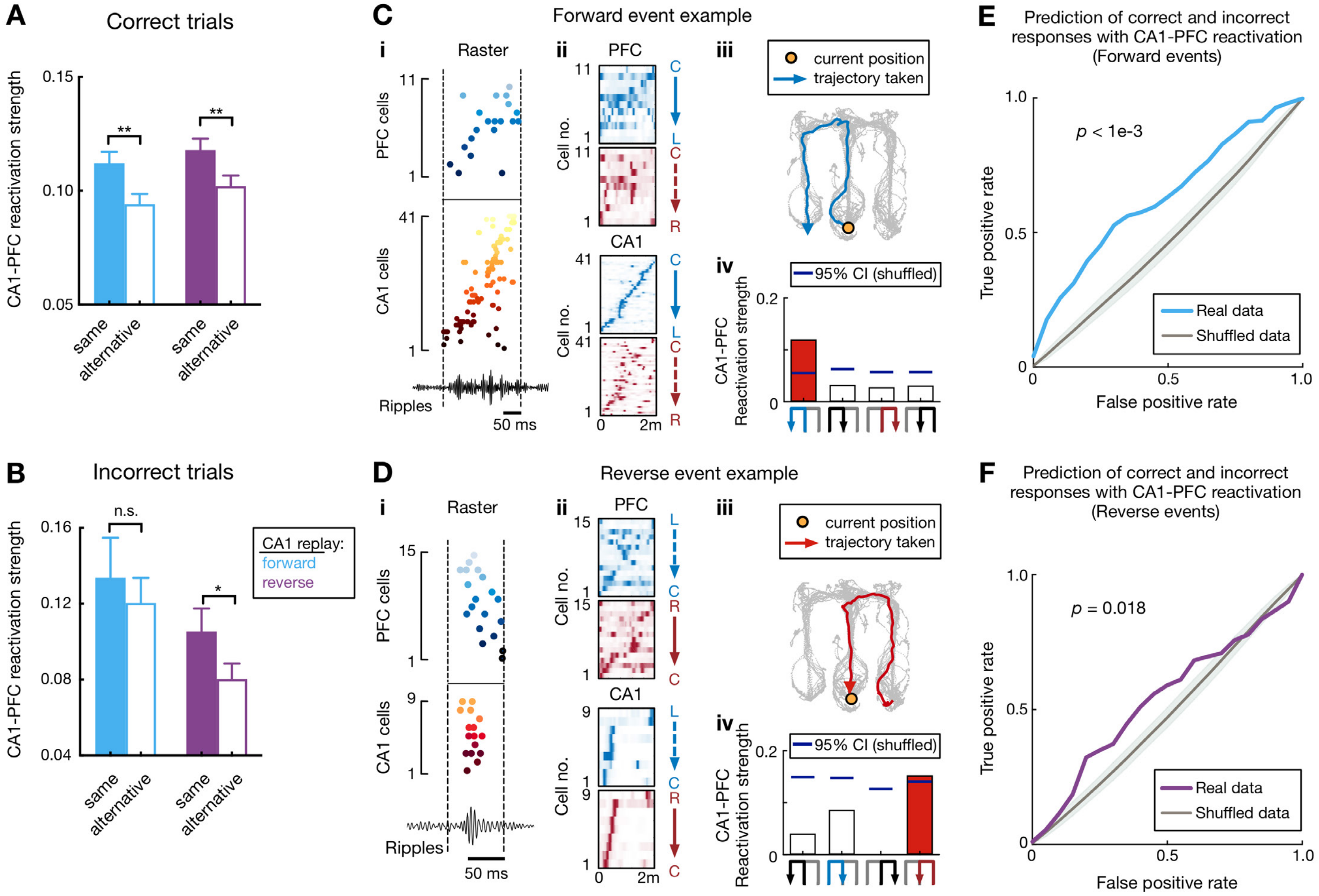


**Figure S7. CA1-PFC reactivation during correct versus incorrect trials**

**(A)** CA1-PFC reactivation of taken path compared to not-taken path during correct (*left*) and incorrect (*right*) trials for forward and reverse CA1 replay (correct trials: **$p = 0.0016$ and $n = 351$ events for forward replay of taken path; **$p = 0.0037$ and $n = 393$ events for reverse replay of taken path; incorrect trials: $p = 0.56$ and $n = 38$ events for forward replay of taken path; *$p = 0.024$ and $n = 48$ events for reverse replay of taken path; event-by-event paired t-tests). Note the robust difference of CA1-PFC reactivation of taken versus not-taken path during correct trials compared to that during incorrect trials. Results were replicated from Figure 7G of Shin et al. 2019 with 3 additional animals. The reactivation strengths of taken (actual) path and not-taken (alternative) path were used as 2D features for the SVM classifiers in **(E)** and **(F)**.

**(C)** Example forward CA1-PFC replay sequences representing actual future choice (see also **Figure 1C** for this event with example cells, and ripples from a different tetrode). **($C_i$)** Ordered raster plot during a

SWR event (black line: ripple-band filtered LFPs from one CA1 tetrode). **($C_{ii}$)** Corresponding spatial fields. **($C_{iii}$)** Actual (immediate future) trajectory (orange circle: current position when replay sequences occurred). **($C_{iv}$)** Reactivation strength (trajectory schematics on the *bottom*). Blue horizontal lines: 95% CIs computed from shuffled data. Red bar: the decoded trajectory.

**(D)** Example reverse CA1-PFC replay sequences representing actual past choice at the center well. Data are presented as in **(C)**.

**(E** and **F)** CA1-PFC replay strength predicts correct and incorrect responses. **(E)** Prediction using replay strength of CA1-PFC forward events. **(F)** Prediction using replay strength of CA1-PFC reverse events. ROC curves were computed for the SVM classifiers (*p*-value from trial-label shuffling denoted; see **METHODS**). Shadings: SDs.
